# A protein standard that emulates homology for the characterization of protein inference algorithms

**DOI:** 10.1101/236471

**Authors:** Matthew The, Fredrik Edfors, Yasset Perez-Riverol, Samuel H. Payne, Michael R. Hoopmann, Magnus Palmblad, Björn Forsström, Lukas Käll

## Abstract

A natural way to benchmark the performance of an analytical experimental setup is to use samples of known content, and see to what degree one can correctly infer the content of such a sample from the data. For shotgun proteomics, one of the inherent problems of interpreting data is that the measured analytes are peptides and not the actual proteins themselves. As some proteins share proteolytic peptides, there might be more than one possible causative set of proteins resulting in a given set of peptides and there is a need for mechanisms that infer proteins from lists of detected peptides. A weakness of commercially available samples of known content is that they consist of proteins that are deliberately selected for producing tryptic peptides that are unique to a single protein. Unfortunately, such samples do not expose any complications in protein inference. For a realistic benchmark of protein inference procedures, there is, therefore, a need for samples of known content where the present proteins share peptides with known absent proteins. Here, we present such a standard, that is based on *E. coli* expressed human protein fragments. To illustrate the usage of this standard, we benchmark a set of different protein inference procedures on the data. We observe that inference procedures excluding shared peptides provide more accurate estimates of errors compared to methods that include information from shared peptides, while still giving a reasonable performance in terms of the number of identified proteins. We also demonstrate that using a sample of known protein content without proteins with shared tryptic peptides can give a false sense of accuracy for many protein inference methods.

## Introduction

Shotgun proteomics offers a straightforward method to analyze the protein content of any biological sample. The method involves proteolytic digestion of the proteins into peptides, which greatly improves the efficiency of the technique, but also introduces a problem for the subsequent data processing. As the mass spectrometers are detecting ions from peptides rather than proteins directly, the evidence for the detection of the peptides has to be integrated into evidence of the presence of proteins in the original sample, using a *protein inference* algorithm [1].

This protein inference procedure is complicated by the homology within most proteomes, many proteins share constituent proteolytic peptides and it is not clear how to best account for such *shared peptides*. For example, should we see shared peptides as evidence for all, a subset, or none of its potential aggregate proteins? While the field of computational proteomics starts to reach consensus on how to estimate the confidence of peptide-spectrum matches (PSMs) and peptides, there is still relatively little work done in establishing standards that evaluate how much confidence we can give reported protein inferences, or even what the best methods to infer proteins from shotgun proteomics data are.

Currently, there are two available methods to determine the accuracy of inference procedures and their error estimates: (i) simulations of proteomics experiments and (ii) analysis of experiments on samples with known protein content. By simulating the proteolytic digestion and the subsequent matching of mass spectra to peptides [2,3,4] one can obtain direct insights into how well the simulated absence or presence of a protein is reflected by a protein inference procedure. However, there is always the risk that the assumptions of the simulations are diverging from the complex nature of a mass spectrometry experiment. Hence, accurate predictions on simulated data can only be viewed as a minimum requirement for a method to be considered accurate [4].

A more direct characterization of protein inference procedures can be obtained by analyzing experiments on samples of known protein content [5]. For such experiments, a protein standard is assembled from a set of isolated and characterized proteins and subsequently analyzed using shotgun proteomics. Normally, the acquired spectra are searched against a database with sequences from proteins known to be present, as well as absent proteins [6]. Two notable protein standards are available today, the ISB18 [5], and the Sigma UPS1/UPS2 standard. Both standards consist of a relatively limited set of proteins, 18 proteins in the ISB18, and 48 proteins in the UPS. However, neither of those standards produce tryptic peptides shared between multiple protein sequences. Hence, these standards are a poor fit for benchmarking protein inference algorithms, as the real difficulty of protein inferences is proteolytic peptides shared between multiple proteins.

Here, we present a benchmark dataset specifically designed for the protein inference problem. Two different samples of defined content were created from pairs of proteins that share peptides. These protein standards are a by-product from the antibody production from the Human Proteome Atlas project (http://www.proteinatlas.org/) [7], where protein fragments, referred to as *Protein Epitope Signature Tags* (PrESTs), are expressed in recombinant *E. coli* strains as antigens to be injected into rabbits to raise polyclonal antibodies. We demonstrate that such protein fragments can be used for benchmark protein inference procedures and for evaluating the accuracy of any confidence estimates such methods produce. This dataset has also previously been made available in anonymized form as a part of the iPRG2016 study (http://iprg2016.org) on protein inference. Here, we also evaluated a set of principles for protein inference with the sets.

## Methods

### Data generation

To generate the datasets, the PrEST sequences of the Human Proteome Atlas-project were scanned for 191 overlapping pairs of PrEST sequences. From these pairs, two pools A and B were created with overlapping peptide sequences. Each pool contained only one of the PrESTs of each pair (Figure 1). A third pool was created by mixing the pool A and B, resulting in a pool A+B. An amount of 1.8 pmol of each PrEST was added to either the pool A or pool B, and an amount of 0.9 pmol of all PrESTs was added to the pool A+B. A total protein amount of 10 *μ*g from each of the pools were reduced with dithiothreitol and alkylated with iodoacetamide prior to trypsin digestion overnight and each pool was mixed into a background of a tryptic digest of 100 ng *Escherichia coli* [BL21(DE3) strain], resulting in three mixtures, Mixture A, Mixture B, and Mixture A+B. In effect, the resulting concentrations of tryptic peptides shared between Mixture A and Mixture B were the same across all three mixtures, while peptides unique to Mixture A or Mixture B appear in half the concentration in Mixture A+B.

**Figure 1:**
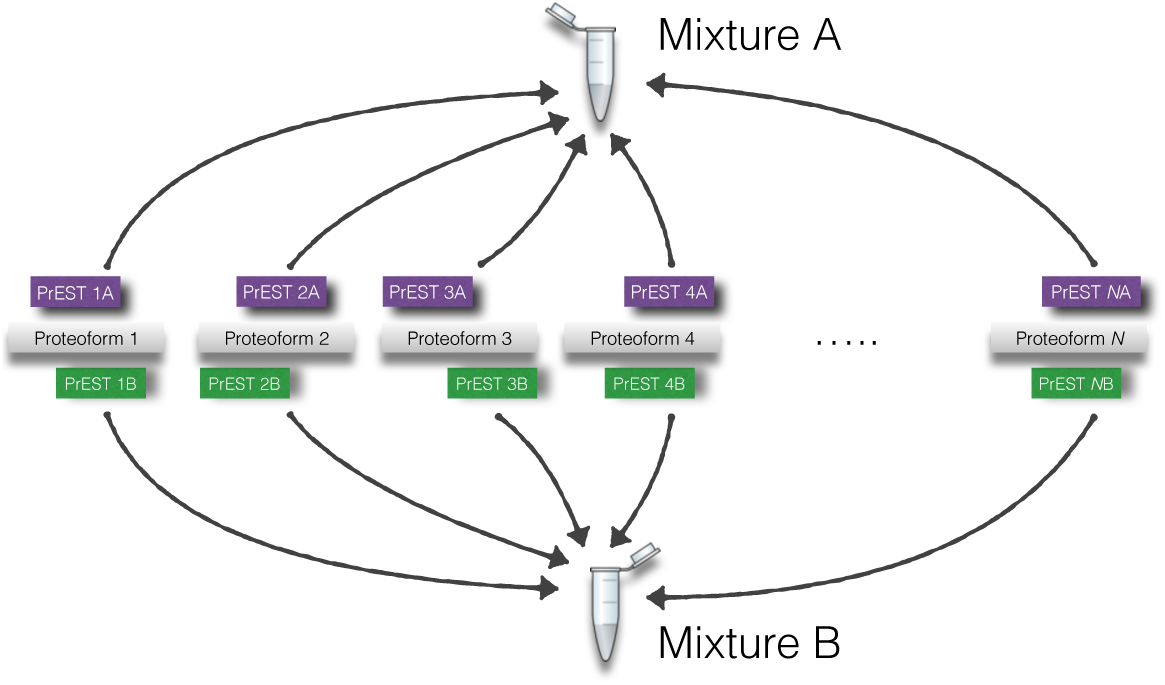
The design of the two mixtures A and B. Two mixtures were generated from 191 overlapping PrEST sequences. We generated mass spectrometry data from the two mixtures individually, as well as from a combination of the two, A+B.

A per sample amount of 1.1 *μ*g of each of three Mixtures was analyzed in triplicate by LC-MS/MS in random order. The digests were loaded onto an Acclaim PepMap 100 trap column (75 *μ*m × 2 cm, C18, 3 *μ*m, 100 Å), washed for 5 minutes at 0.25 *μ*L/min with mobile phase A [95% H_2_O, 5% DMSO, 0.1% formic acid (FA)] and thereafter separated using a PepMap 803 C18 column (50 cm × 75 *μ*m, 2 *μ*m, 100 Å) directly connected to a Thermo Scientific Q-Exactive HF mass spectrometer. The gradient went from 3% mobile phase B [90% acetonitrile (ACN), 5% H_2_O, 5% DMSO, 0.1% FA] to 8% B in 3 min, followed by an increase up to 30% B in 78 minutes, thereafter an increase to 43% B in 10 min followed by a steep increase to 99% B in 7 min at a flow rate of 0.25 *μ*L/min. Data were acquired in data-dependent (DDA) mode, with each MS survey scan followed by five MS/MS HCD scans (AGC target 3e6, max fill time 150 ms, mass window of 1.2 *m*/*z* units, the normalized collision energy setting stepped from 30 to 24 to 18 regardless of charge state), with 30 s dynamic exclusion. Both MS and MS/MS were acquired in profile mode in the Orbitrap, with a resolution of 60,000 for MS, and 30,000 for MS/MS.

### Dataset

We assembled the data into a test dataset consisting of:

- Three FASTA-files, containing the amino acid sequence of the Protein fragments of Mixture A, Mixture B, as well as the sequences of 1000 representative protein fragments that are known to be absent from the sample.
- Twelve runs, consisting of triplicates of analyses of Mixture A, Mixture B, Mixture A+B and “blank” runs without spike-ins, all run in a background of *E. coli*-lysates. These are provided in Thermo raw data-format.
- An evaluation script, written in python.

The FASTA-files and scripts can be downloaded from https://github.com/statisticalbiotechnology/proteoform-standard, and the mass spectrometry data is accessible from the pride database under the project accession number PXD008425.

This dataset has been made available in anonymized form as a part of the iPRG2016 study (http://iprg2016.org) on protein inference. The sample mixtures can be made available on request, for evaluation under other mass spectrometers than the one we used in this study.

### Data Processing

The raw data files were converted to MS1 and MS2 format using ProteoWizard [8] and subsequently processed by Hardklor [9] followed by Bullseye [10], through the interface of the Crux 2.1 package [11], to set monoisotopic masses. The resulting ms2 spectra were then matched to separate target and decoy sequence databases (described below) using Crux 2.1 [12, 11] and Percolator v3.01 [13, 14], deriving peptide-level probabilities for each of the mass spectrometry runs.

The target protein database consisted of all three FASTA-files combined and the decoy database was constructed by reversing the protein sequences of the target database.

For each of the runs we calculated protein-level *Entrapment FDRs* [6] by counting all matches to PrESTs present in their analyzed mixture as correctly matched, and all matches to PrESTs absent (i.e. stemming from the set of 1000 non-present PrESTs or from the set of the mixtures not used in the sample) as being incorrectly matched. As the correct matches only map to present proteins, whereas incorrect matches distribute over both present and absent proteins, we also normalized the Entrapment FDR by the prior probability of the PrEST to be absent, the so-called π_*A*_ [4].

## Results

We constructed two mixtures, each containing 191 PrESTs, that is one out of each of the 191 pairs of PrEST sequences with partially overlapping amino acid sequence (Figure 1). We analyzed the two samples as well as a combination of the two using LC-MS/MS (see the Methods section).

The three datasets make an informative benchmark set. By matching the spectra of the dataset against a bipartite database containing both the present and some absent PrEST sequences we obtain a direct way to count the number of inferred PSMs, peptides, and proteins stemming from non-present PrESTs [6]. More specifically this allows us to assess the fraction of identifications in a set that stems from absent PrESTs, which we here will refer to as the *Entrapment FDR.* However, unlike traditional samples of known content, this standard contains overlapping protein fragments, which allows us to assess the performance of protein inference algorithms in the presence of homology.

### Protein inference

We tested a set of different protein inference algorithms against our test set. We first analyzed the data using Crux [12, 11] and Percolator [13, 14], deriving peptide-level probabilities for each combination of triplicate mass spectrometry runs. We subsequently compared the performance of the different schemes for inferring proteins and their confidence.

First, there are different ways to infer proteins from peptide sequences. The major difference between the methods relates to how they handle so-called *shared peptides*, that is peptide sequences that due to homology or other reason could stem from more than one protein. The tested inference methods were:

Inclusion – Possibly the easiest way to handle shared peptides is to assign any found peptide to all its possible causative proteins. Under this assumption, we infer the presence of any protein which links to an identified peptide.
Exclusion – Another method is to remove any shared peptides before any reconstruction takes place. Under this assumption, we infer the presence of any protein which links to an identified peptide unique to the protein.
Parsimony – A method that is quite popular for handling shared peptides is to use the principle of parsimony, i.e. to find the minimal set of proteins that would best explain the observations of the PSMs with a score above a given threshold. This principle has been implemented in a couple of well-known software tools such as IDPicker 2.0 [15], and MaxQuant [16]. In cases where there are multiple such minimal sets of proteins, several strategies can be used: do not include any of the sets, apply some form of protein grouping (see the Discussion section), or select one of the sets, either at random or based on the order the proteins are listed in the database. Here, we have opted to use the latter alternative, to select one of the sets at random.

### Methods to rank protein identifications

More than just inferring the protein sequences, any practically usable protein inference strategy has to assign confidence estimates in terms of posterior probabilities or false discovery rates. One way to assign such protein-level statistics is by investigating decoy ratios, a process that depends on assigning scores to rank our confidence in the different protein sequences. The different methods to obtain protein-level scores differ in the way they combine the confidence estimates of the proteins’ constituent peptides. We tested five different methods to score proteins:

**Products of PEPs** – This method summarizes a score for the protein’s constituent peptides by calculating the product of peptide-level posterior error probabilities (PEPs) [16]. This method has been extensively used by tools such as MaxQuant [16], PIA [17] and IDPicker [15].
**Fisher’s method** – Another method that relies on an assumption of independence between different peptides’ incorrect assignment to a protein, is Fisher’s method for combining independent *p* values, which is a classical technique for combining *p* values [18]. Fisher’s method takes into account all constituent peptides of a protein [19, 20, 21], by summarizing the individual peptides empirical *p* values. Unlike the product of PEPs, which also combines peptide-level evidence, Fisher’s method explicitly accounts for the number of *p* values being combined and hence normalizes for protein length to some extent.
**Best peptide** – Instead of weighting together peptide-level evidence for a protein, some investigators chose to just use the best available evidence for a protein [22]. Savitski *et al*. [23] showed that, on large-scale data sets, taking the best-scoring peptide as the representative of a protein was superior to incorporating information from lower-scoring peptides. This approach might feel unsatisfying for most investigators, as the method discards all information but the best-scoring PSM for each protein.
**Two peptides** A simple way to combine evidence at the peptide level is the widely used two-peptide rule [24]. This approach requires evidence for a second peptide to support a protein inference, thereby preventing so-called “one-hit wonders”, *i.e*., cases where a single, potentially spurious PSM yields a spurious protein detection [22]. Furthermore, the recently published Human Proteome Project Guidelines for Mass Spectrometry Data Interpretation version 2.1 requires “two non-nested, uniquely mapping (proteotypic) peptides of at least 9 aa in length”, to count a protein sequence as being validated with mass spectrometry [25].
**Fido** A more elaborate method to estimate the confidence in inferred proteins is to use Bayesian methods, represented here by Fido [26], which calculates posterior probabilities of proteins’ presence or absence status given the probabilities of peptides correctly being identified from the mass spectrometry data. Such methods are normally seen as inference procedures, but due to the design of our study, we listed it as a confidence estimation procedure to be used in combination with Inclusion inferences. Fido’s use requires selection of prior probabilities for protein presence, present proteins emitting detectable peptides and mismatches. These probabilities were set using grid searches, as implemented when running Fido through Percolator [13].

In Supplementary Table S3 we have mapped a set of commonly used protein inference tools according to the names of the inference and ranking principles we use in this paper.

### Performance

First, we measured the performance of the permutations of inference and confidence estimation procedures, in terms of the number of identified proteins at a 5% protein-level entrapment-FDR (see Table 1 and S1, as well as Figure S1). It is worth noting that there is a fundamental difference between the different benchmark sets, for the set A+B all proteins that share tryptic peptides are present. However, the sets A and B, contain tryptic peptides shared between absent and present proteins.

**Table 1:** **The number of inferred proteins at a 5% protein-level entrapment FDR from the peptides derived from the triplicate runs.** Note that, since we are inferring proteins known to be present, any incorrect inferences will be added as additional proteins. Hence the maximal number of inferred proteins is 5% higher than the number of proteins in the mixtures.

| Inference Principle | Scoring method | A | A+B |
| --- | --- | --- | --- |
| Anticipated number of PrESTs | | $191 \cdot 1.05 = 201$ | $382 \cdot 1.05 = 401$ |
| Inclusion | Fisher's method | 112 | 390 |
|  | Products of PEPs | 124 | 395 |
|  | Best peptide | 0 | 395 |
|  | Two peptides | 0 | 381 |
|  | Fido | 181 | 388 |
| Exclusion | Fisher's method | 182 | 345 |
|  | Products of PEPs | 184 | 355 |
|  | Best peptide | 185 | 355 |
|  | Two peptides | 171 | 309 |
| Parsimony | Fisher's method | 181 | 365 |
|  | Products of PEPs | 181 | 365 |
|  | Best peptide | 174 | 365 |
|  | Two peptides | 181 | 344 |

Comparing the different methods of dealing with shared peptides, we found that the Inclusion and Parsimony methods reported more proteins than the Exclusion method when investigating the A+B set that contains peptides shared between present proteins. However, for the A set that contains peptides shared between the present and absent proteins, the Exclusion reports more proteins than the Inclusion methods (except Fido) and the Parsimony methods.

For the different confidence estimation procedures, we noted that the TWO PEPTIDES method reported fewer proteins than the other methods, whereas FIDO reports more proteins than the other methods.

Many implementations of Parsimony are two-step procedures, which first threshold on peptide-or PSM-level FDR and subsequently infer the most parsimonious set of proteins. In such implementations, one ends up controlling the list of proteins at both peptide and protein-level. We chose to make a more extensive test series of Parsimony for 1%, 5% and 10% peptide-level FDR (See Figure S3 and Table S2). A trend is observable for such data, for the A+B set, more proteins are observed for similar protein-level FDRs when using a higher peptide-level FDR (e.g. 10%) than when using a lower peptide-level FDR (e.g. 1%). However, the inverse is true for the A set, more proteins are observed for similar protein-level FDRs when using a 1% peptide-level FDR than when using a 10% peptide-level FDR. This is a consequence of the cases where the algorithm has to select one of many minimal sized subsets explaining the observed peptides at random. Such a selection is not harmful to the performance if all minimal subsets consist of present proteins, but might be harmful if one or more of the minimal subsets contains absent proteins [27].

### Accuracy of confidence estimates

Subsequently, we set out to estimate the accuracy of the different confidence estimates of the protein inference procedures and hence plotted the entrapment FDR as a function of the methods’ reported FDR in Figure 2 and Supplementary Figure S2.

**Figure 2:**
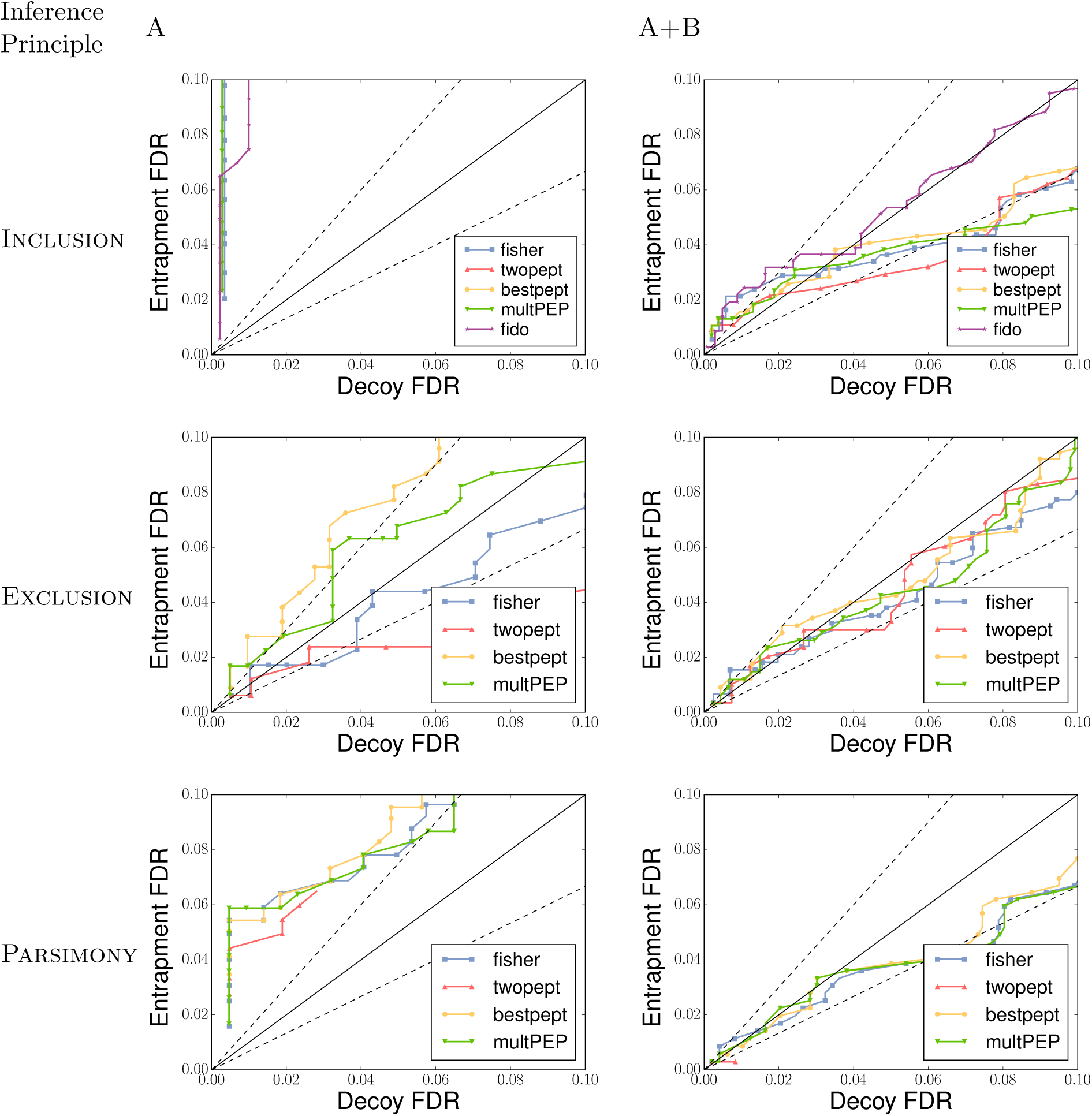
The accuracy of the tested confidence estimation procedures for different inference methods. The figures plots reported *q* values from the decoy model, the decoy FDR, against the fraction of entrapment proteins in the set of identified target proteins, the observed entrapment FDR using a peptide-level FDR threshold of 5%. The dashed lines indicate *y* = *x*/1.5 and *y* = 1.5*x*.

All the tested methods reported acceptable and similar accurate statistics for the sets where the shared peptides stem from proteins that are all present (set A+B). However, when comparing the decoy and entrapment FDR for the sets with peptides shared between present and absent proteins (set A) we see that none of the inference methods using Inclusion or Parsimony reported satisfying decoy-derived FDRs.

The Exclusion principle, on the other hand, seemed to handle the sets with peptides shared between present and absent proteins in a satisfying manner, and particularly the Fisher’s method and the Products of PEPs gave quite accurate statistics for such data without reducing the number of reported proteins.

## Discussion

Here we have described a protein standard that can be used for comparing different algorithmic approaches to inferring (sorted) lists of proteins from shotgun proteomics data. The set is particularly useful for determining how to handle protein inferences in cases where a peptide could stem from multiple different proteoforms. We used the dataset to compare a set of approaches for protein inference and protein scoring models, and found that the reliability of protein inferences became more accurate when excluding any peptides shared between multiple proteins as compared to, *e.g*. inferring the most parsimonious set of proteins.

Many algorithms and tools that use Parsimony or Inclusion principles, group proteins according to the peptide evidence that they share. Particularly in the case of Inclusion, the accuracy would increase dramatically if we would have evaluated protein groups rather than individual proteins. We have not included this option here for the sake of simplicity, not least because it confounds the null hypothesis in a way that complicates a fair comparison between methods [27, 4].

While the set is larger and more complex than other samples of known content (several millions of spectra), we see room for future improvements both in terms of more and longer protein sequences than our current standard as well as more complicated patterns of shared peptides.

Recently, a couple of other benchmarks for protein inference algorithms have been published. First, when selecting a protein inference strategy for Percolator, the authors used simulations to show that excluding shared peptides and scoring the protein based on the best scoring peptide performed overall better than the compared methods. However, on small-sized datasets, the method of multiplying PEPs had a slight performance advantage [14].

Second, a large center study, Audain *et al*. [28], benchmarked a set of protein inference algorithm implementations on “gold standards” i.e. manually annotated datasets. The authors conclude that PIA [17] and Fido [26] perform better than the other analyzed implementations on their datasets. We did not include PIA in our comparisons, but we did include Fido. In line with Audain *et al*., Fido gave an excellent performance and calibration and on datasets where all proteins sharing a peptide were present (set A+B). However, Fido’s assessment of reliability scores was less than ideal for the datasets where the shared peptides’ causative proteins were from both the present and absent groups (set A and set B). This behavior could not have been characterized using datasets lacking peptides shared between multiple proteins, and it is hence not a surprise that such characteristics have not been noted in previous studies.

The two-peptide rule performed poorly, as has been reported by several studies [22, 29, 30]. For instance, Veenstra *et al.* wrote already in 2004 that “Simply disregarding every protein identified by a single peptide is not warranted” [22]. Similarly, the FDRs reported after using parsimony to handle shared peptides seems, in general, to be anti-conservative which is in line with what was reported by Serang et al., “parsimony and protein grouping may actually lower the reproducibility and interpretability of protein identifications.” [27].

In this study we have kept the peptide inference pipeline identical for all protein inference pipelines, to enable a *ceteris paribus* comparison of the protein inference methods. We also have tried to benchmark principles rather than implementations but made an exception for the Fido inference method, as this was readily available in the Percolator package that was used for the peptide inferences.

## Acknowledgments

S.H.P. was supported by the US Department of Energy, Office of Science, Office of Biological and Environmental Research, and Early Career Research Program. M.R.H. was supported by the National Institutes of Health, National Institute for General Medical Sciences (grant R01 GM087221) and Center for Systems Biology (grant 2P50 GM076547) and the National Science Foundation MRI (grant 0923536). L.K. was supported by a grant from the Swedish Research Council (grant 2017-04030).

